# Diversity and population genetic structure of the wax palm *Ceroxylon quindiuense* in the Colombian Coffee Region

**DOI:** 10.1101/443960

**Authors:** Natalia González-Rivillas, Adriana Bohórquez, Janeth Patricia Gutierrez, Víctor Hugo García-Merchán

**Affiliations:** Grupo de Investigación en Evolución, Ecología y Conservación (EECO), Programa de Biología, Universidad del Quindío, Carrera 15 Calle 12 Norte, Armenia, Quindío, Colombia; Grupo de Investigación y Asesoría en Estadística, Universidad del Quindío; International Center for Tropical Agriculture (CIAT), Km 17, recta Cali-Palmira, Colombia

**Author notes:** (NGR), (AB), (JPG) & (VHGM). The authors mentioned contributed equally to this work.

**Keywords:** Neotropical Palm, *Ceroxylon quindiuense*, Genetic variability, Microsatellites

## Abstract

The wax palm from Quindío (*Ceroxylon quindiuense*) is an icon of the cultural identity of the coffee growing eco-region and of all Colombia. Processes of urbanization, expansion of the agricultural and livestock area, among others, have increased its level of threat. Protecting this palm from extinction is important at an ecological level, given its function as a key species in Andean ecosystems. This work evaluated the diversity and population genetic structure of the wax palm from Quindío in five populations of the Colombian coffee region eco-region (Andean zone) by using ten microsatellite molecular markers. Two groups were identified at genetic diversity level, along with a heterozygote deficiency in all the populations possibly due to cryptic population structure effects mediated by loss of habitat. The five sampling units considered presented a total significant genetic structure, revealing a high degree of reproductive isolation. The results presented here add to the Conservation Plan for this species existing in Colombia.

## Introduction

Loss of habitat is the processes with the highest impact on tropical ecosystems [1]. This process has obvious effects, like modification of the landscape and local elimination of some species; but can also have effects on the long-term viability of populations of certain species due to the reduction of the number and isolation of its individuals [2]. From a genetic point of view, diminished effective population size and increased degree of isolation makes the fragmented populations more susceptible to genetic drift and inbreeding, diminishing success and the evolutionary potential of the species and increasing the risk of extinction [3, 4].

The wax palm from Quindío (*Ceroxylon quindiuense*) is the Colombian national tree [5]; it is distributed in the country’s three mountain ranges of the Andes [6], often being a dominant plant from the montane rain forest canopy and considered a key species within the ecosystems it inhabits [7]. It is a fundamental part of the structure and diversity that makes up the Colombian coffee cultural landscape, playing a crucial role as source of food for wildlife, like birds and small mammals [7, 8].

This species is currently in danger of extinction (EN), according to the red list of palm species [9]. Some of the threats affecting *C. quindiuense* and, generally, palms of the genus include their commercial use through extraction of leaves for the Catholic Holy Week, an activity that has been managed and reduced considerably [9]. Other threats include paddocking processes which involves the loss of habitat and low regeneration because of the introduction of other uses of the soil they occupy (mainly crops and livestock). This considerable reduction of its habitat has permitted estimating that their population has diminished by over 50% in the last three generations (210 years) [10].

Bearing in mind the latent threats this species has for its survival, it becomes relevant to know the effect caused by the loss of habitat on the persistence, genetic diversity, and future viability of umbrella species, like *C. quindiuense*, which currently has no defined conservation areas in Colombia. Thus, knowledge and understanding of the diversity and genetic structure of a species plays an important role for the conservation and analysis of ecological and social aspects; permitting among some points, to determine the response capacity of the populations and species against natural environmental changes or those provoked by conscious or unconscious human activities [11]. Likewise, the study of the genetic constitution of the populations and of the mechanisms of evolutionary change acting on them are the pillars that constitute the tools to confront the loss of biological diversity [12].

For this reason, it is important to diagnose the current situation and conservation of the wax palm from Quindío, focusing on the genetic level of distribution in the studied areas. This diagnosis will permit undertaking actions to protect this palm from extinction, adding that the main problem identified, the loss of biological diversity of the species, is the impact that can be generated in the continuity of the landscape and other species that depend on it. Likewise, *C quindiuense* is the flagship species of the eco-region of the Colombian coffee region, a zone declared world heritage by UNESCO in 2011, which generates a big opportunity to continue with conservation efforts of the Colombian national tree.

In genetic structure studies on *Ceroxylon*, Gaitán [13] investigated the genetic diversity and population structure in three palm species (*C. sasaime*, *C. alpinum,* and *Attalea amigdalina*) immersed in a fragmented landscape in the central Andes of Colombia. Evaluations of genetic diversity using microsatellite molecular markers in *C. sasaimae* and *C. alpinum* showed that the populations conserve high genetic diversity in spite of their restricted distribution, loss of habitat, and the low number of surviving individuals. Additionally, no significant differences were found in genetic diversity among seedlings, juveniles, and adults in any of the species studied.

Sanín [14] explored two aspects simultaneously, phylogeography and conservation, from microsatellites of nine populations of *C. quindiuense*, *C. ceriferum,* and *C. ventricosum*, describing their diversity gradients within the context of the tropical Andes. The epicenter of the genetic diversity of the *C. quindiuense* is in the Colombian central mountain range, given that both populations studied there gathered most of the alleles and the broadest allelic range. Likewise, a highly significant genetic structure was noted among all the populations, a high degree of reproductive isolation, and the almost absence of potentially mixed individuals, suggesting no recent gene flow in said populations.

Chacón-Vargas [15] compared the genetic diversity and population structure of three *Ceroxylon quindiuense* nurseries evaluated with that of the Colombian sylvan populations from Valle de Cocora in the Department of Quindío and Toche and La Línea in the Department of Tolima. Use of microsatellite molecular markers of *C. sasaimae* and *C. alpinum* developed by Gaitán [13] demonstrated that the populations of this species in the central mountain range have a high genetic diversity that permits establishing potentially successful repopulation programs. This also exhibits significant genetic structuring, suggesting that we should seek the conservation of each unit that represents an independent gene pool.

Considering the studies mentioned, it is expected that populations of *Ceroxylon quindiuense* in the Colombian coffee eco-region will be also highly structured with limited gene flow among them. This would generate patterns of low genetic diversity and, with such, a high degree of inbreeding as a consequence of the loss of habitat present for this species. Due to all the aforementioned, this study evaluated the diversity and population genetic structure of the wax palm from Quindío (*Ceroxylon quindiuense:* Ceroxyloideae) in five populations of the Colombian coffee eco-region by using microsatellite molecular markers.

## Materials and Methods

### Study area

The area of the Colombian coffee region represents a set of ecosystems distributed between the central and western cordilleras of the Colombian Andes, where ecological and human complexes coexist, showing indivisibility comprised by the unit of basins with slopes and plains. This includes the snow-capped mountains; that border the biogeographic Chocó; the coffee region ecosystem, and the complex urban corridor [16], which previously only permitted designating the departments of Caldas, Quindío, and Risaralda. Now, it is used to identify a territory that, besides these three departments, covers the north of the department of Valle del Cauca and the northwest of Tolima [16]. This work included five locations distributed in the departments of Caldas, Quindío, Tolima, and Valle del Cauca (Fig 1).

**Fig 1.**
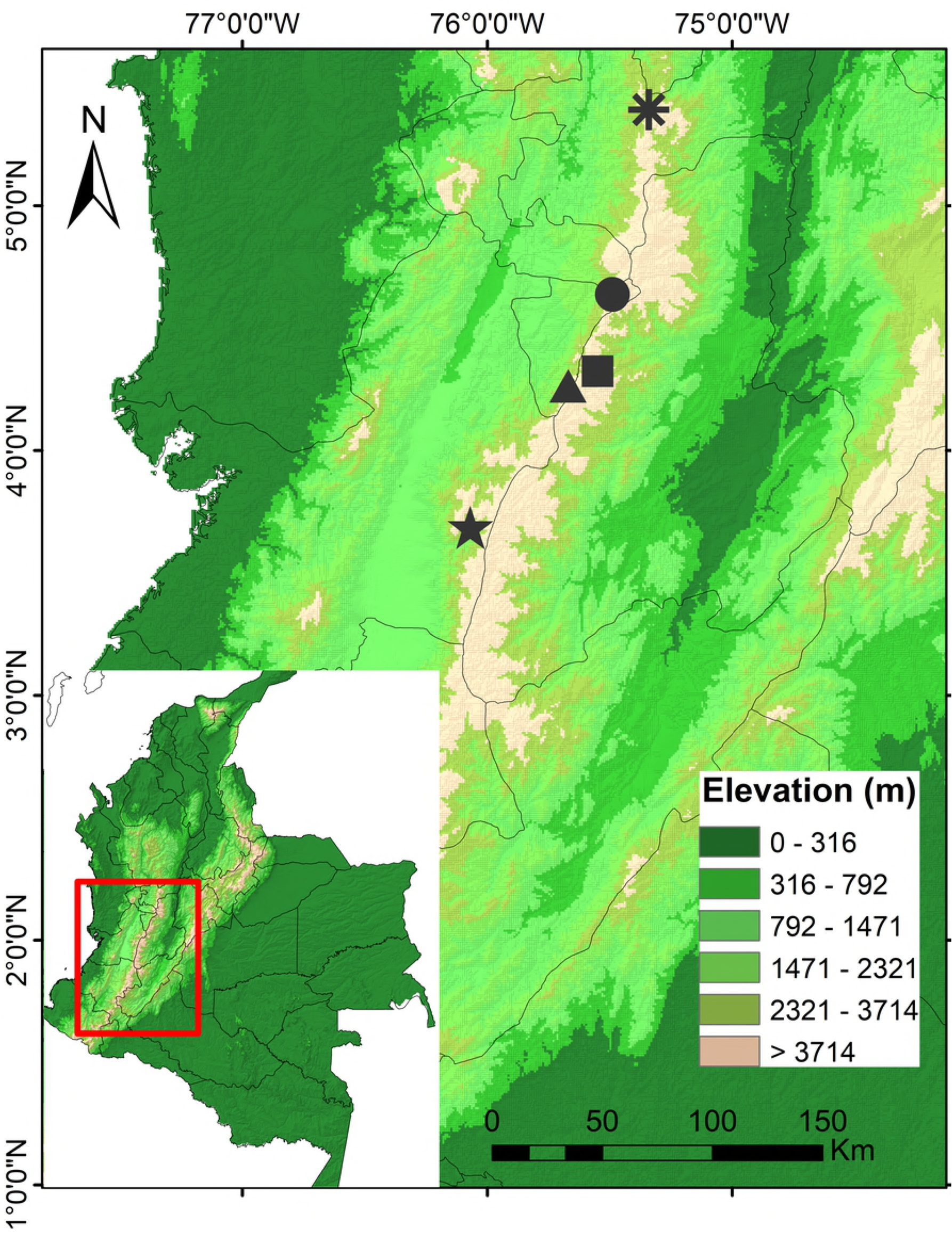
Locations for sampling of the wax palm from Quindío (*Ceroxylon quindiuense)* in the coffee area eco-region: Circle (Valle de Cocora, Salento, Quindío, 2300-2900 masl); Square (Anaime, Cajamarca, Tolima. 2000-3000 masl); Star (Tenerife, Valle del Cauca. 2500-2600 masl); Asterisk (San Félix, Salamina, Caldas. 2800 masl); Triangle (Pijao, Quindío. 2600-3000 masl)

### Sampling design and collection of plant material

Ten individuals were collected for each of the three development stages, seedlings, juveniles – less than 10 m high – and adults – with prior corroboration of the reproductive state of *C. quindiuense* (Fig 2) in wooded areas with different degrees of intervention, defined as the size of each of the patches chosen. For each of the individuals sampled, the corresponding coordinates were taken with the aid of an Etrex global positioning system. The study avoided collecting nearby individuals to discard bias in the diversity analysis caused by kinship under the closest neighbor model, taking 100 m as the minimum distance between individuals.

**Fig 2.**
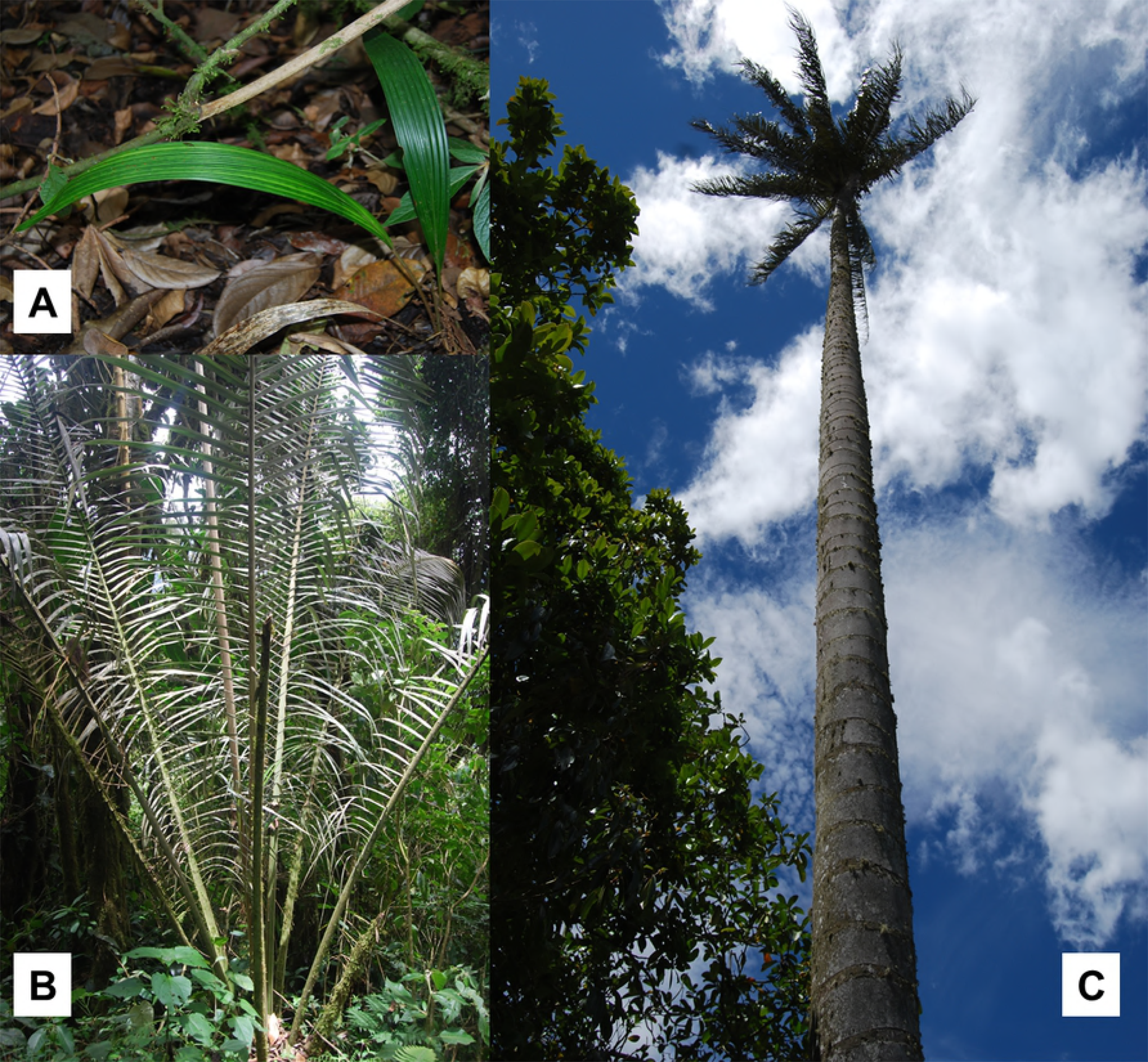
Representatives of each of the three development stages of wax palm from Quindío (*Ceroxylon quindiuense*) sampled. **A.** Seedling, **B.** Juvenile, **C.** Adult

The plant material collected in the three development stages were tender leaves stored in tightly sealed bags containing silica gel in 1:5 proportion (tissue: silica) to accelerate dehydration of the tissues. Collection of the adult individuals was conducted by scaling to the crowns of the palms and collecting (inasmuch as possible) the tenderest leaves from the adults. Certified personnel carried out this activity with all the stipulated security measures. All the procedures carried out in the field and access to the genetic resource followed the guidelines defined in the resolution by the entity responsible for it, *Corporation Autónoma Regional from Quindío* (CRQ), No. 1136 of December 31, 2012 and Decrees 1376 and 1375 of the Ministry of the Environment and Sustainable Development, both from 27 June 2013, respectively.

### DNA Extraction

The DNA was extracted by taking 1g of previously macerated tissue from the leaf collected and using the Qiagen DNeasy Plant Mini kit (Qiagen, Ltd.), following the protocol proposed by the manufacturer. The concentration of the material extracted in ng/μL was determined by using NanoDrop 1000 Spectrophotometer V3. The DNA visualization was carried out through agarose gel at 1% stained with the SYBR^®^ Safe colorant by Invitrogen, using 3 μl of DNA and 2 μl of Blue Juice. Thereafter, dilutions were made of said material at a concentration of 8 ng/ml at a final volume of 50 μl and stored at −20 °C.

### Evaluation and amplification of loci microsatellites

Ten loci microsatellites (SSR) were evaluated, which were developed by Gaitán [13] for the species of the genus *Ceroxylon alpinum* and *C. sasaime* (Table 1).

**Table 1.**
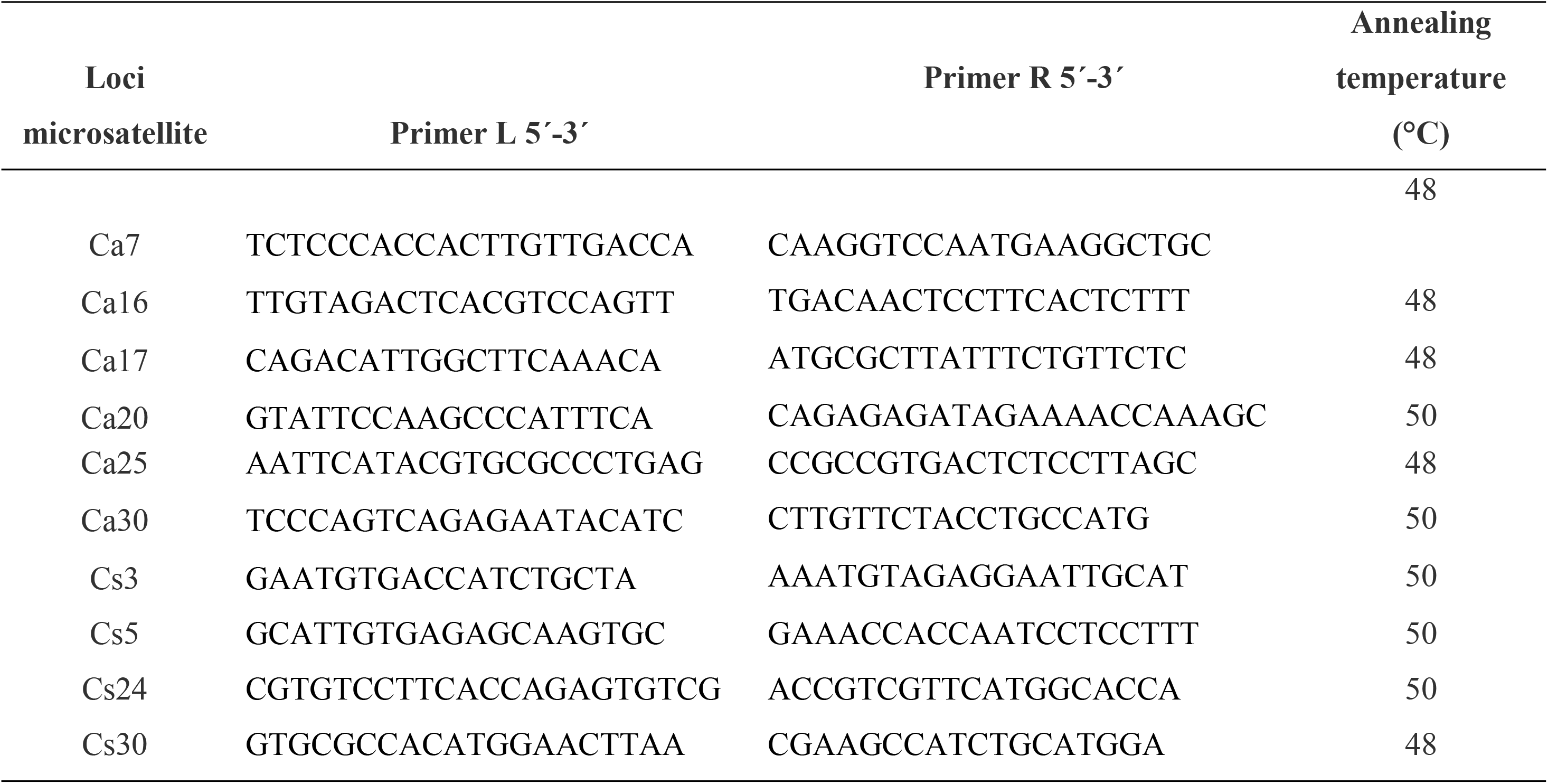
Loci microsatellites used and annealing temperature of each evaluated in five populations of wax palm from Quindío (*Ceroxylon quindiuense*).

Each loci used was diluted by using 10 μl of the primer and 90 μl of sterile water. Each amplification reaction through PCR used 25 μl of final volume, consisting of: 5 μl of DNA (8 ng/μl), 12.5 μl of GoTaq^®^ Colorless Master Mix by Promega Ltd, 1 μl of primer L-R and 5.5 μl of water free of nucleases by Promega Ltd. The DNA and the GoTaq^®^ Colorless Master Mix were kept in ice when out of the refrigerator to avoid degradation at ambient temperature. Amplifications were carried out in a PTC-100 thermocycler (MJ Research) using the following amplification program: 94 °C (4 min), [94 °C (15 s), annealing temperature 46-50 °C (15 s), 72 °C (5 min)] for 35 cycles, 72 °C (5 min), 10 °C (∞). The subsequent visualization of the PCR products was conducted in agarose gels at 3% (5 μl PCR product + 3 μl Blue Juice).

### Amplification of fluorescent microsatellites

Amplification of the microsatellites for their subsequent analysis was via the automated fragments analysis methodology, based on the detection of fluorescent colorants that permits rapid selection of polymorphic sequences to multiple genomic loci [17]. To reduce the cost of genotyping with microsatellites marked with fluorescence, Oetting *et al.* [18] proposed a PCR strategy called multiplex with primers or tail primers, which employs a *forward* primer with an additional extension of 18 pb in its 5′ extreme that belongs to the M13 primer (TGTAAAACGACGGCCAGT), a *reverse* primer without altering, and a third universal M13 primer marked with fluorescence.

Amplifications for the microsatellites with end labeling were carried out on 96-well plates. The cocktail for PCR used per reaction was conducted at a final volume of 20 μl, mixing 5 μl of DNA (8 ng/μl); 1X buffer; dNTP 0.3 mM; magnesium chloride (MgCl_2_) 3.5 Mm; 0.1 pmol/μl of the M13 primers marked with fluoro-chrome and *reverse*; 0.05 pmol/μl of the *forward* primer with the M13 sequence; and 0.5 U of *Taq pol.* The quality of the products amplified was verified in agarose gels at a concentration of 1.5%, taking 5 μl of product and 3 μl of Blue Juice.

Upon obtaining all the amplification products, the mixture was made according to the size range in base pairs of the microsatellites. These mixtures were carried out on 96-well skirt plates, which combined the PCR product of the primers selected for each of the combinations taking a 2-μl aliquot of each product amplified to be sent to Cornell University (Ithaca, NY. USA) and its subsequent analysis through the Applied Biosystem 3730xl DNA sequencer (Applied Biosystem).

### Data analysis

The size of the alleles, in base pairs, was estimated through the GeneMarker software v.2.6.7 (SoftGenetics, State College, PA). A matrix of individuals and microsatellites was generated, indicating the diploid genotypes for each marker. To test the adequate assignment of the genotypes, the AlleloBin program (ICRISAT) was used for each. This program tested if the sizes of fragments observed adjusted to the size of alleles expected, based on the units of length repeated in the microsatellite [19].

### Structure and diversity analysis

The allele frequencies, the observed heterozygosity (*H_O_*) and expected (*H_E_*) per locus and per population were calculated by using the GeneAlEx 6.41 software [20]. Deviations from the Hardy-Weinberg Equilibrium (HWE) and the linkage disequilibrium were evaluated for each locus using the GENEPOP software v. 4.1 [21], employing Markov’s chain method with 5000 iterations and following the algorithm by Guo and Thompson [22].

Genetic divergence among populations was estimated by calculating the classical approach of the *F_ST_* and the likelihood ratio logarithm (G test) to evaluate population differentiation through Markov chain algorithms, as implemented in the GENEPOP software [23, 24]. Although *F_ST_* has been widely used as a measure of population structure, it has recently come under criticism due to its dependence on the intra-population heterozygosity [25]. Hence, the population differentiation was also estimated by using the *D_ST_* statistics, which more precisely reflects the genetic differentiation among populations because it includes a multiplicative partition of the diversity, based more on the number of alleles than on the heterozygosity expected [25]. For its calculation, the DEMEtics package [26] was used from R v.3.3.1 [26], with a bootstrap method of 1000 repetitions to estimate the level of significance [27]. An analysis of isolation by distance was performed through a Mantel test [28], correlating the geographic and genetic distances (*F_ST_* and *D_ST_*) using the GenAlex 6.41 software [20].

The identification of the number of genetically differentiated groups was carried out using the Bayesian clustering method [29] implemented in STRUCTURE, version 2.3.3, and the discriminant analysis of principal components (DAPC: [31]) available in the adegenet package [30] for R, version 3.3.1 [26]. Both methods do not require *a priori* delineation of genetic clusters and are suitable for analyzing spatially continuous data [29, 31]. However, some differences exist in the analytical approaches between the two methods. STRUCTURE attempts to cluster individuals by minimizing Hardy–Weinberg and gametic disequilibrium [29], and typically fails to identify some complex types of spatial structure such as isolation-by-distance [31] and hierarchical population structure [32]. The multivariate analysis used in DAPC does not make any assumption on the population genetic models and may be more efficient at identifying genetic clines and hierarchical structure [31]. The two dissimilar approaches were used in this study, because various clustering approaches may lead to different conclusions [e.g. 33–35].

For DAPC, the data were scaled and all the data lost (absent) were replaced with the value of the median (origin of the X and Y axes). A number of principal components (PCs) were considered as predictors for the DA. Nevertheless, no strict criterion exists to determine how many PCs/DAs must be considered during this dimension reduction step, beyond considering the statistical power that implies having the highest possible amount of PCs/DAs and the stability of the assignments carried out [31]. This study considered 42 PCs and 2 DA, containing 80% of the total genetic information. For this analysis, the populations genotyped were used *a priori* to assign the genotypes. The discriminant functions and the genotypes of the microsatellites were visualized as diagrams created with the *scatter.dapc* function in the R “adegenet” package [30, 31].

Likewise, an analysis based on a Bayesian model implemented in software STRUCTURE v. 2.3.3 [29] was used to estimate the number of groups (K) represented by all sampled individuals and individuals mixture proportions. The number of groups was inferred using 10 independent simulations with Markov chains 100000 (MCMC), following the model that allows the origin of individuals from mixed populations with correlated allele frequencies and using the locality as *a priori* information to help identify groups. The range of K groups to be analyzed for between 1 and 10. The most likely K was estimated using Evanno’s DK [32], which was implemented in Structure Harvester [36].

## Results

### Genetic variability of *Ceroxylon quindiuense*

In the five locations sampled (Cocora, Anaime, Pijao, San Félix, and Tenerife), 237 alleles were identified in the 10 loci studied for 126 individuals. The range of alleles per locus varied from 11 to 35. The Ca17 and Cs03 loci presented the highest number of alleles observed for all the locations (35 and 30, respectively). Locus Ca07 presented the lowest number of alleles observed (11). In Cocora, the minimum number of alleles was seven (locus Ca07); six in Anaime (locus Ca07); four in Pijao (locus Ca16); five in San Félix (locus Cs24) and five in Tenerife (locus Ca30). The number of alleles observed in Cocora was 117; 101 in Anaime; 95 in Pijao; 116 for San Félix and 119 in Tenerife (Fig 3).

**Fig 3.**
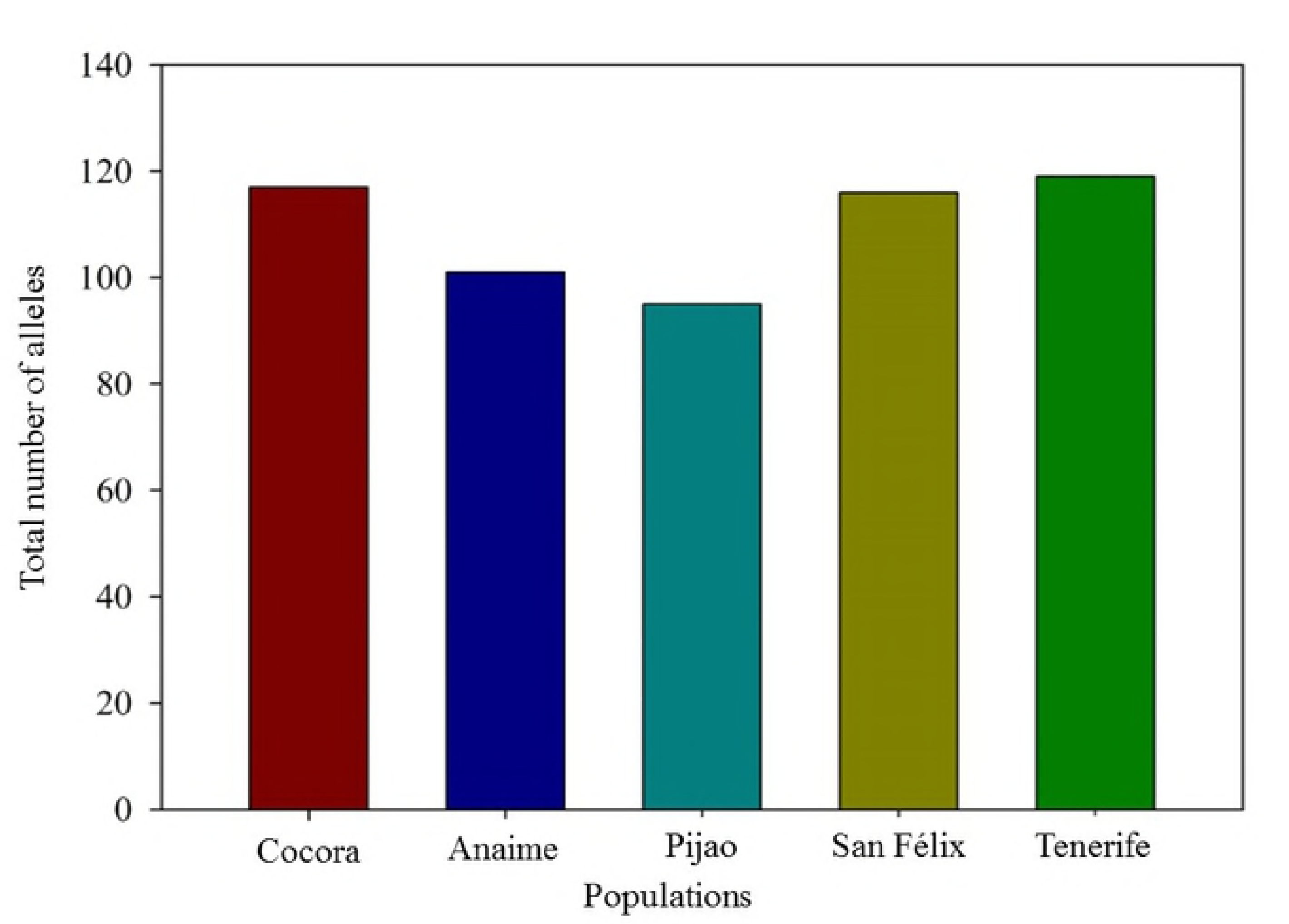
Number of alleles for the 10 loci microsatellites evaluated in five populations of wax palm from Quindío (*Ceroxylon quindiuense*)

In general, the allele frequencies for the 10 microsatellites presented a similar distribution with frequencies between 10 and 30%; however, frequencies above 50% were identified in four locations (Anaime, Pijao, San Félix, and Tenerife) and frequencies below 42% in the population of Cocora. Allele 142 in loci Ca30 and Ca20 presented allele frequencies above 50% in the locations of Anaime and San Félix. Likewise, allele 216 showed an allele frequency above 50% for locus Cs05 in the location of Pijao. The highest allele frequency registered was evidenced for allele 117 and 145 of the loci Ca30 and Ca25, respectively, in Tenerife.

### Observed heterozygosity and expected, effective number of alleles (*Ne*), HWE deviations, and private alleles

In genetic diversity, two groups were found: San Félix, Anaime, and Pijao comprised the first with an observed heterozygosity (*Ho*) of 0.343 and Cocora and Tenerife comprised the second with *Ho* of 0.216. The expected heterozygosity (*He*) fluctuated between 0.739 (Tenerife) and 0.828 (Anaime/Cocora). The fixation index (*F*) varied between 0.509 (San Félix) and 0.772 (Tenerife) (Table 2). A total number of homozygote genotypes of 256 and 196, respectively, was registered for Tenerife and Cocora (populations with higher *F*). The 10 loci evaluated were 100% polymorphic. The effective number of alleles (*Ne*), defined, as the required number of alleles with equal allele frequency to predict the heterozygosity, was determined for each locus in all the locations evaluated. The *Ne* fluctuated between 6.572 (San Félix) and 7.247 (Cocora).

**Table 2.** Number of individuals samples (n), mean values of fixation index (*F*), observed heterozygosity (*Ho*), expected heterozygosity (*He*), and number of alleles for the 10 loci evaluated in five populations of wax palm from Quindío (*Ceroxylon quindiuense*)

| Population | n | F | Ho | He | Number of alleles |
| --- | --- | --- | --- | --- | --- |
| San Félix | 30 | 0.509 | 0.390 | 0.803 | 116 |
| Anaime | 20 | 0.609 | 0.310 | 0.828 | 101 |
| Cocora | 26 | 0.704 | 0.283 | 0.828 | 117 |
| Pijao | 20 | 0.573 | 0.330 | 0.804 | 95 |
| Tenerife | 30 | 0.772 | 0.150 | 0.769 | 119 |

Globally, the value of the inbreeding coefficient (*F_IS_*) was positive and highly significant (p ≤ 0.001) for all the populations evaluated when considering all the loci, indicating that said populations presented a defect of heterozygotes and, hence, are not in Hardy-Weinberg Equilibrium. The value of *F_IS_* was 1 for locus Ca25 in four populations (Cocora, Anaime, Pijao, and San Félix), for locus Ca17 in Anaime, and loci Ca07 and Cs30 in Tenerife, indicating that there are only homozygote individuals for these loci.

The private alleles (unique) were 17 for Cocora, 16 for Anaime and Pijao, 21 for San Félix, and 35 for Tenerife (Fig 4). For the populations of Cocora and Pijao, locus Ca17 showed the highest value with five private alleles, in Anaime loci Ca17 and Ca20 with four, in San Félix locus Ca25 with six, and for Tenerife loci Cs03 and Ca20 with 12 private alleles recognized (Fig 5).

**Fig 4.**
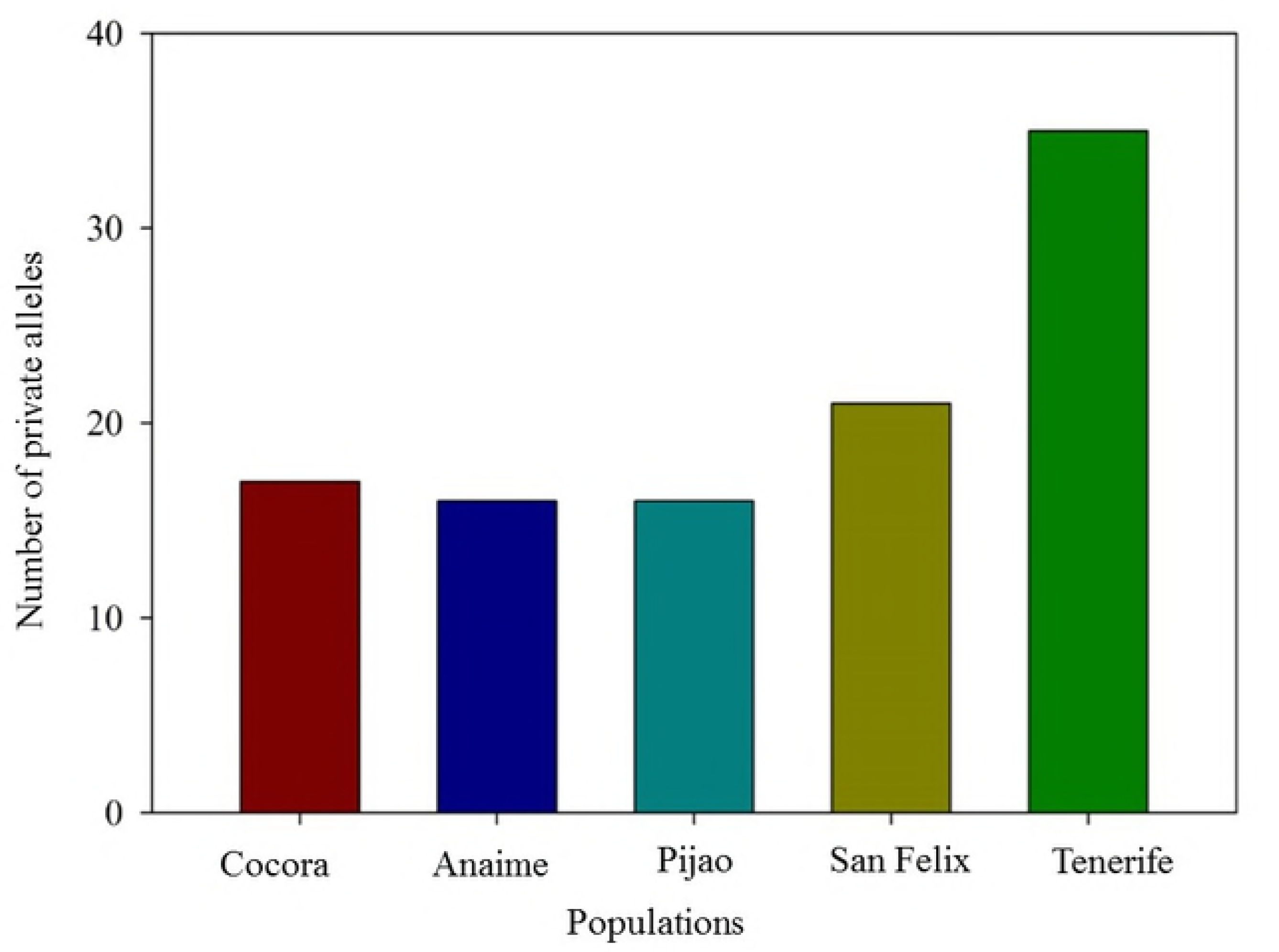
Total number of private alleles (unique) for five populations of wax palm from Quindío (*Ceroxylon quindiuense)*

**Fig 5.**
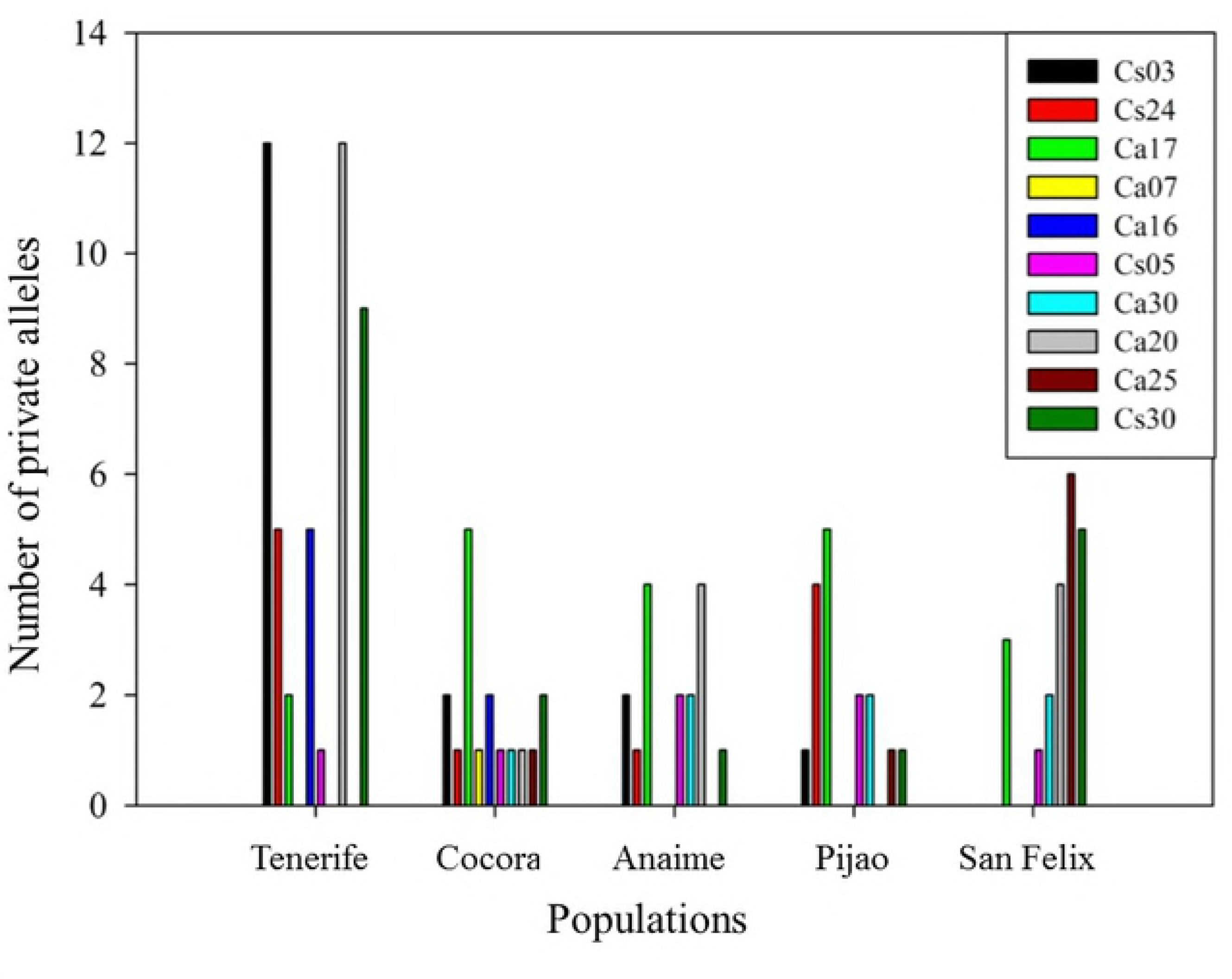
Total number of private alleles (unique) for each of the 10 loci evaluated in five populations of wax palm from Quindío (*Ceroxylon quindiuense*)

### Genetic population structure of *Ceroxylon quindiuense*

The five sampling units considered in this study had a total significant genetic structure and by pairs of populations. The population of Tenerife showed the highest values of differentiation for both structure indices (*F_ST_* and *D_ST_*) with respect to the rest. The populations of Pijao and Anaime had the lowest values for both indexes; nonetheless, they were also significant (Table 3). Analysis of seedlings and juveniles for all the populations presented no significant differences within each population (p > 0.05).

**Table 3.**
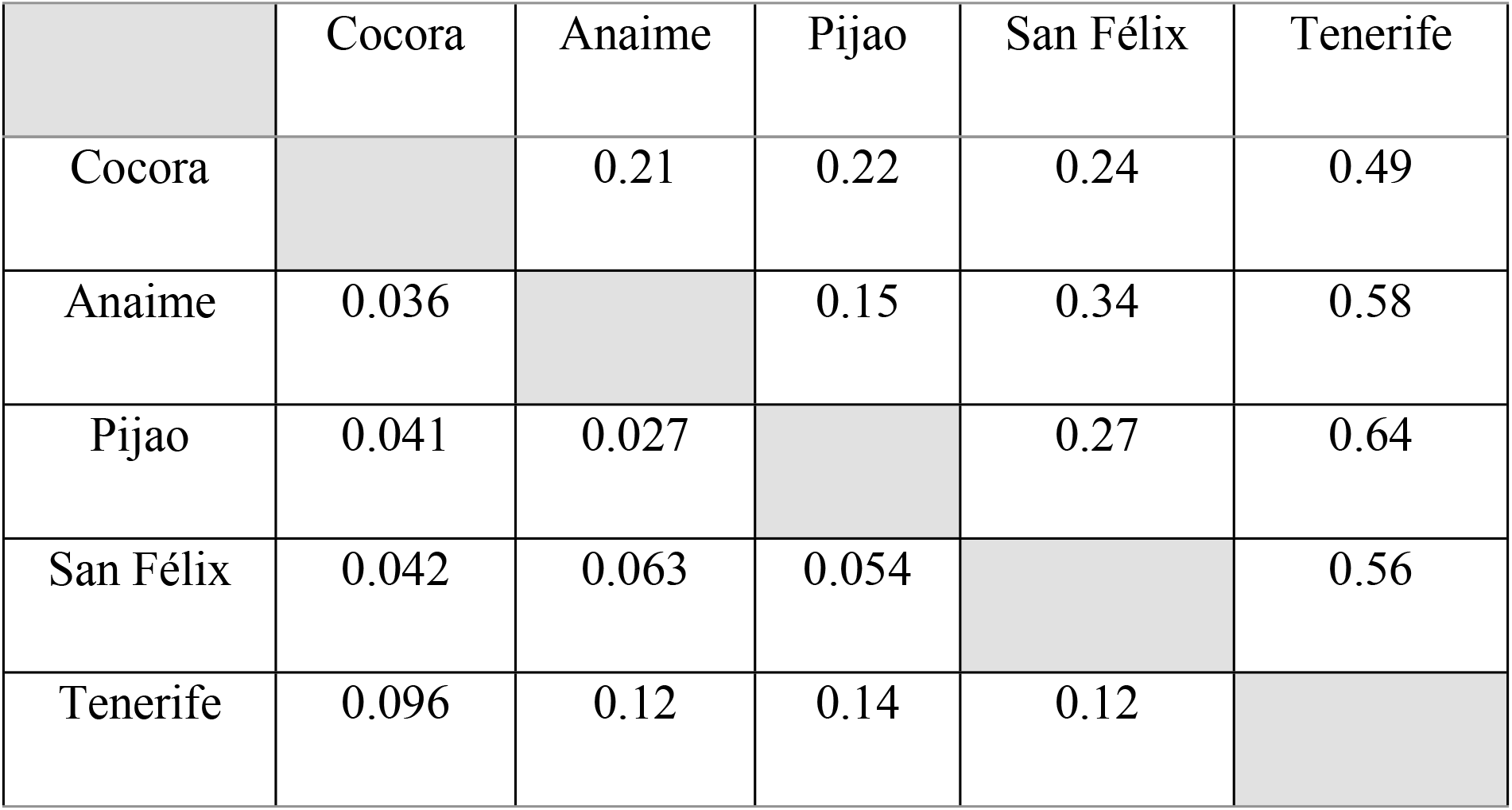
Genetic structure indexes among the five populations of wax palm from Quindío (*Ceroxylon quindiuense*). *F_ST_* below the diagonal (p < 0.05) and *D_ST_* above the diagonal (p < 0.001).

The analysis of isolation by distance through the Mantel test was not significant for *D_ST_* or for *F_ST_* vs. geographic distances. The DAPC permitted identifying three genetic groups in which, nonetheless, a mixture of individuals is found from different populations, except for a group that presents in its majority individuals from the population of Tenerife (Fig 6). For other hand, Bayesian analysis using STRUCTURE identified two cluster (K=2), reflecting a small mix of individuals between clusters (Fig 7).

**Fig 6.**
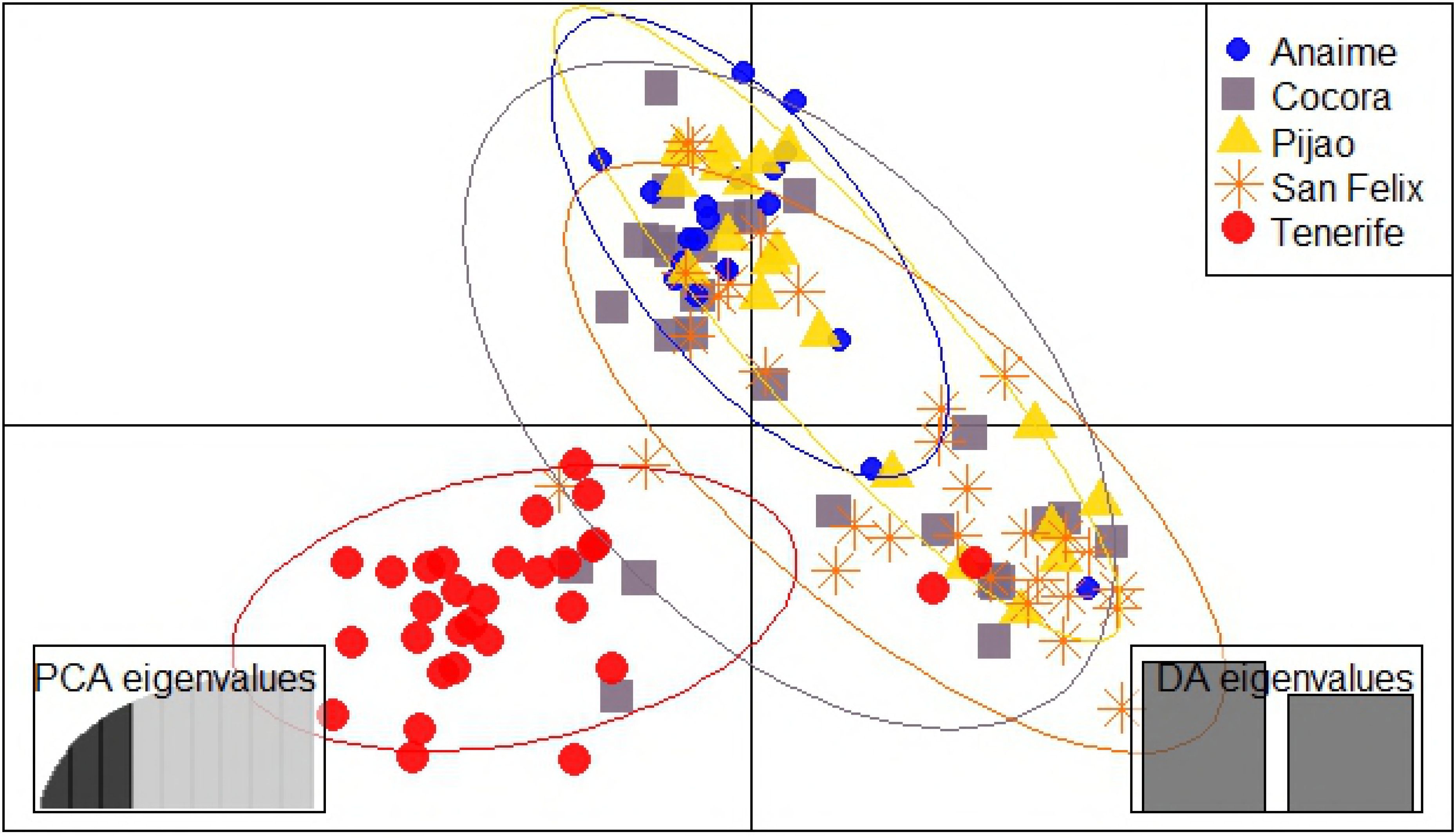
Discriminant analysis of principal components with three groups *(k-means)*. Genetic profile based on 10 microsatellites, 126 individuals collected in five populations of wax palm from Quindío. Each individual per population is represented by a specific symbol and color. Ellipses only permit showing the differentiation of the three groupings identified

**Fig 7.**
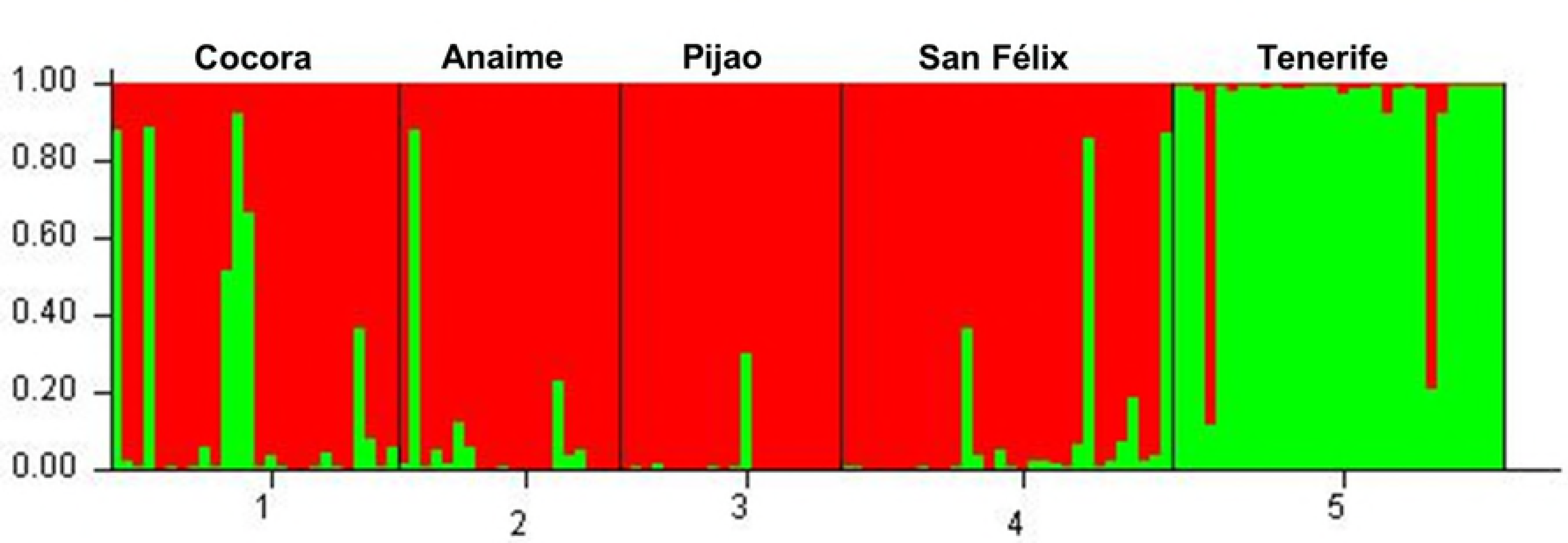
Graphical representation of the genetic composition of all individuals evaluated. Analysis of group membership for all individuals. The colors represent the genetic groups, such that the color ratio for each individual indicates their probability of assignment to the genetic cluster

## Discussion

In this study, the wax palm from Quindío and Colombian national tree, *Ceroxylon quindiuense*, showed variable patterns of genetic diversity and significant genetic differentiation at structure level in the natural populations evaluated from the eco-region in Colombia’s coffee growing area. Two groupings were identified in genetic diversity. The first, constituted by the populations of Cocora and Tenerife, characterized for presenting low diversity with lower values of observed heterozygosity (*Ho*) and high fixation index (*F*); the second was made up of the populations of Anaime, Pijao, and San Félix, exhibiting high diversity and higher *Ho*, and a lower value of *F* (Table 2).

Previous studies have registered that diversity in terms of observed heterozygosity for *C. quindiuense* populations, such as Cocora, has been higher than that identified in this study (*Ho* = 0.467), as well as in other populations of this species, like Tolima (*Ho* = 0.51), Antioquia (*Ho* = 0.536), and Santander (*Ho* = 0.450) [37, 15]. Features such as allelic range, the effective number of alleles *(Ne),* and heterozygosity values suggest a high genetic diversity for populations of wax palm from Quindío, a pattern explained as a result of the long-life cycle of the species and its ability to survive for more than 120 years after deforestation [10], in addition to the sexual system that is present the species.

However, a high percentage of populations with heterozygote deficiency has been determined for this species [14] and for others from the genus *Ceroxylon*, such as *C. alpinum*, *C. sasaime* [13], and *C. equinulatum* [38]. This agrees with the inbreeding coefficient values (*F_IS_*) determined in this study, thus, identifying heterozygote deficiency in all of the populations studied.

In the natural populations, heterozygote deficiency is generally attributed to aspects like selection of loci, selection against heterozygotes, and inbreeding [39]. Nevertheless, for *C. quindiuense* it is assumed to be generated through a phenomenon known as cryptic population structure [14], where the gene flow in a subdivided population is hindered by the existence of geographic barriers, habitat fragmentation, among others; with fragmentation being one of the principal threats to which all the populations of this species are subjected. Galeano et al. [9] state in the Conservation Plan for said species, that a considerable portion of the individuals that exist survive in paddocks. A large proportion of individuals is found in relatively small forest fragments, whose long-term permanence is not guaranteed.

The impact of loss of habitat due to human activities is reflected on the structure and on the amount of genetic variation of the populations [40], as observed in this study and in other studies developed for this species, identifying a high degree of inbreeding. Seed dispersion is an important process for the dynamics of the populations because it combines demographic and genetic effects [41]. A scenario of dispersion limitation due to fragmentation may lead to diminishing the effective number of parentals at a brief temporal scale and increasing the genetic drift in successive generations, especially when dominant mothers exist with a high gametic contribution (*i.e*., correlated maternity) or events of correlated dispersion [42, 43].

Dispersion through grains of pollen is also a mechanism that is a fundamental part in the gene flow of plant species; fragmentation of the habitat has also been postulated as a negative effect on populations of wild pollinators, but to date, relatively few studies exist on the effects of fragmentation on pollination by itself. Pollination in *C. quindiuense* is conducted by several species of nitidulid beetles of the tribe Mystropini [44], which consume floral parts and pollen. They masticate while remaining in the inflorescence; part of the pollen is adhered to their bodies and they then transfer it to another inflorescence.

The relationship between the wax palm from Quindío and said beetles, as well as the effect of fragmentation in this process, are aspects still not studied. However, it has been recorded that many species of pollinators, like the nitidulidae, would have sufficient flight capacity [45–47]; to be able to displace between sites crossing the matrix that surrounds the fragments and, thus, exploit the floral resources available in the different forest sites. It has been shown that local populations of pollinators would have the capacity to survive and remain in each site, independent of their flight capacity and, hence, would not be greatly affected by the diminished area [48]. Effective dispersion mechanisms at medium and long distance can permit relatively high levels of gene flow in fragmented systems, while restricted dispersion systems, or those failed over time and/or space, can cause interruption of the genetic connectivity [49]. Although the pollen dispersion (haploid) at long distance is translated into an important genetic connectivity mechanism, the limitation in seed dispersion (diploid) implies limitation in gene distribution – both maternal and paternal [41].

*C. quindiuense* has a strong genetic structure with all the populations differentiated, yielding significant structuring values (Table 3); results agreeing with previous studies in populations, like Cocora (Quindío), Roncesvalles, Toche, and la Línea (Tolima), Jardín (Antioquia) and Zapatoca (Santander) [14,15]. This fact reveals a high degree of reproductive isolation for the wax palm populations. Said degree of reproductive isolation identified between these populations is not associated directly to phenomena of isolation by distance. Nevertheless, Sanín et al. [37] state that *C. quindiuense* populations, as well as those of other species of the genus, are genetically isolated as result of their phylogeographic history occurring throughout the formation of the Andes and their migration processes.

Even so, it is evident that the current state of conservation of its habitat is influenced considerably in said structuring, where landscape characteristics may modify gene flow between population pairs, directly when affecting the dispersion rates between them, or indirectly upon affecting the spatial disposition of the dispersion rates and among the subpopulations intervened [50]. For *C. quindiuense*, the isolation by resistance (IBR) model can be applied [50]. It is a method designed to predict the effects of the landscape’s structure on the equilibrium of the genetic structuring in natural populations, incorporating the landscape’s heterogeneity within the analysis of gene flow and of genetic differentiation evaluated for this species.

Landscapes associated to the wax palm from Quindío are characterized for undergoing diverse transformation phenomena that has turned them into heterogeneous landscapes, reporting that the populations from the nation’s central mountain range have been deforested principally during the 1900s, during a process of landscape transformation called “Paisa Colonization” [37]. This has given way to the current existence of a landscape with forest fragments of diverse sizes, large areas of paddocks and/or of diverse crops. Using the IBR model might permit predicting the patterns of movement and the probabilities of successful dispersion of *C. quindiuense* through complex landscapes, like those presented, to generate connectivity measures, define isolation of the patches or populations, and identify important connective elements (for example, corridors) to plan their conservation [51].

Although prior studies have reported the near absence of individuals potentially mixed in this species suggesting no recent gene flow [14], the DAPC conducted in this study demonstrates the presence of mixed individuals among the three groupings identified. This aspect is possibly related to the fact that both slopes of the central mountain range have served as migrant receptors and, hence, it is where the gene flow has historically converged from the north and south of the species distribution [37]. This result highlights the importance of evaluating three components that principally make up the gene flow, namely: a spatial component (distance), a bioclimatic component (habitat), and a biological component, as studied in *C. echinulatum* [38] that permit determining a possible gene flow dynamic that would give way to increased genetic variation among the populations.

It was identified that the population of Tenerife had the highest number of private alleles, the lowest levels of genetic diversity (Table 2), and significant structuring with respect to the other populations (Table 3); even the majority of individuals formed one of the groupings identified in the DAPC (Figure 6) and the STRUCTURE analysis (Figure 7). This population has not been studied previously; however, its low genetic diversity and population genetic structure may be attributed to historical and contemporary processes of colonization (i.e. expansion of the agricultural, livestock area and processes of urbanization), which has led to the isolation, as it has been reported by Sanin et al. [37] in other populations of *Ceroxylon*.

The genetic structure of a population is determined by its evolutionary history and expresses the amount of genetic diversity it shelters and how it is distributed within the population [52]. It was possible to determine that the significant structuring levels from Tenerife, as named, cannot be attributed to the considerable geographic distance where the other populations studied are found, proving through the analysis performed of isolation by distance. In addition to current processes, the inbreeding phenomenon that has occurred with the other populations was evaluated. With this being an almost completely deforested zone, with livestock and crops of scallions, banana passion fruit (curuba), blackberry, and potato, etc., which makes it the first agricultural pantry in Valle del Cauca, containing fragments of wasteland and very disturbed forest [10]. Thus, it may be inferred that this population has undergone processes at local scale, like the lack of dispersers that distribute the seeds within the forest, thereby generating the need to conduct studies to assess the behavior of the dispersers after destruction and fragmentation events of this and all the populations of *C. quindiuense.*

The genetic criteria obtained from the study of molecular markers could be useful to select wildlife populations as candidates with priority for their conservation [53]. As the case of the populations of wax palm from Tolima and Quindío for which Sanín [14] states that the huge genetic diversity available in these populations, in spite of the current loss of habitat, may serve this species as source to repopulate in case of an eventuality that causes local extinctions in the satellite populations. However, it is important to remember that the small populations or satellites of wax palm from Quindío, as well as Tenerife, present important distinguishable genetic patterns such as their low observed heterozygosity, their high values of homozygotes and fixation index, and their higher number of private alleles. Finally, we recommend using methods based on phylogeography and molecular phylogenetics that help to reconstruct historical patterns that have led to the current geographic distribution [54–56]. In said scenario, the wax palm from Quindío represents an excellent model to carry out these types of studies that explain processes of historical species diversification in the Andes.

This study is, to date, the first work at genetic level developed for the populations of San Félix, Anaime, Tenerife, and Pijao that contributes to one of the first lines of action of the Conservation Plan for this Andean palm in Colombia, broadening the knowledge base in this species. Thus, we recommend considering the genetic diversity identified for said satellite populations and associating them to the other populations studied to generate a complete diagnosis that permits its conservation on both sides of the central mountain range, as suggested in the Conservation Plan. Likewise, these results will permit complementing the conservation actions proposed in this document, such as the preservation and restoration of its habitat, with greater focus on the interconnection among forest fragments and populations through the renovation of palm groves in the paddocks and forested areas. This should lead to increasing its distribution area and diminishing vulnerability to local extinction of satellite populations; adding aspects related to protection of the local dispersers that play an important role in gene flow.

## Acknowledgments

This research was funded by project 673 of the Vice-rectory of Research at Universidad del Quindío. Gratitude is also expressed to the International Center for Tropical Agriculture and especially to Dr. Joe Thome and to Johan Carvajal and Jessie Ortíz for their support in the fieldwork. We also thank Alicia Knudson for improving the English of this manuscript.

